# Temperature drives seagrass recovery across the Western North Atlantic

**DOI:** 10.1101/2024.07.31.605761

**Authors:** Fee O. H. Smulders, Justin E. Campbell, Andrew H. Altieri, Anna R. Armitage, Elisabeth S. Bakker, Savanna C. Barry, S. Tatiana Becker, Enrique Bethel, James G. Douglass, Hannah J. van Duijnhoven, Jimmy de Fouw, Thomas K. Frazer, Rachael Glazner, Janelle A. Goeke, Gerrit Gort, Kenneth L. Heck, Olivier A. A. Kramer, Ingrid A. van de Leemput, Sarah A. Manuel, Charles W. Martin, Isis G. Martinez López, Ashley M. McDonald, Calvin J. Munson, Owen R. O’Shea, Valerie J. Paul, Laura K. Reynolds, O. Kennedy Rhoades, Lucia M. Rodriguez Bravo, Amanda Sang, Yvonne Sawall, Khalil Smith, Jamie E. Thompson, Brigitta van Tussenbroek, William L. Wied, Marjolijn J. A. Christianen

## Abstract

Climate-driven shifts in herbivores, temperature and nutrient runoff threaten coastal ecosystem resilience. However, our understanding of ecological resilience, particularly for foundation species, remains limited due to a rarity of field experiments that are conducted across appropriate spatial and temporal scales and that investigate multiple stressors. This study aimed to evaluate the resilience of a widespread tropical marine plant (turtlegrass) to disturbances across its geographic range and how this is impacted by environmental gradients in (a)biotic factors. We assessed the resilience (i.e. recovery) of turtlegrass to a simulated disturbance (complete above- and belowground biomass removal) over a year. Contrary to temperate studies, higher temperature generally enhanced seagrass recovery. While nutrients and light availability had minimal impact, combined high levels of nutrients and herbivore grazing (meso and megaherbivore) reduced aboveground recovery. Our results suggest that the resilience of some tropical species, especially in cooler subtropical waters, may initially increase with warming.

## Introduction

Understanding the ability of coastal ecosystems to resist and recover from disturbances, i.e. ecosystem resilience^1^, is essential in an era of rapid global change; yet we know very little about the impact of anthropogenic stressors on the resilience of key foundation species that cover our coastal zones. This is cause for concern as global warming and other human-induced stressors are increasingly driving large-scale ecosystem loss^2^, and coastal ecosystems are among the most threatened^3,4^. Often empirical studies sample and compare resilience at a single location and focus on the effects of single stressors on aboveground traits. This approach leads to an incomplete understanding of the impact of environmental drivers on ecosystem resilience across large spatial scales, and a lack of practical resilience indicators to aid management in future global change scenarios.

The resilience of coastal plant communities is under pressure from both global change stressors such as warming seas and intensifying storms, as well as human pressure from population-dense coastal zones^5,6^. Deterioration of coastal ecosystems will cause a concurrent loss of ecosystem services^7^, such as efficient carbon storage capacity^8^ and coastal protection^9^. Because of their thermal tolerance limits, population range shifts and mass mortality of coastal foundation species are expected under projected global change scenarios^10,11^. Furthermore, warming increases the frequency and intensity of heatwaves and storms^12,13^, which may lead to declines^14^. Meanwhile, climate-induced poleward shifts of herbivores can lead to local changes in grazing pressure that can negatively impact coastal plants^15,16^. Additionally, local impacts include eutrophication due to urban and agricultural nutrient runoff^17^, which can cause loss through algal proliferation that results in light limitation in aquatic systems^18^. These factors that are often assessed singularly may interact and cause vegetation decline by compromising stability and thus ecological resilience^6,19^. For example, seagrasses weakened by eutrophication may be more vulnerable to heat stress^20^, increasing the chance of meadow collapse.

Field-based methods testing the resilience of foundation species are rapidly developing. In particular, increasing evidence suggests that dynamic indicators, such as the recovery rate after a disturbance, may better indicate the resilience of an ecosystem compared to static indicators, such as cover or standing biomass^21,22^. It is rarely feasible to apply system-wide perturbations to experimentally assess resilience in vegetated habitats. Instead, measuring the recovery rate after a small-scale experimental perturbation can serve as a reliable indicator of the resilience of a large-scale ecosystem^23,24^, where a slow recovery after a physical disturbance may signal ecosystem vulnerability to a phase shift^19^. Disturbance and recovery experiments – mimicking pulse disturbances such as storms^25^ – thereby provide a tool to determine the overall resilience of an ecosystem. In macrophyte dominated marine ecosystems, the focus is often on measuring aboveground recovery^26,27^. However, knowledge of belowground dynamics is key for understanding the resilience of these ecosystems^28,29^, because belowground biomass includes the carbon reserves important for recovery potential^16,30^ and the root structure that provides stability and resistance to uprooting by waves and storms^31,32^. Therefore, both belowground dynamics and dynamic indicators such as recovery rates are essential to include in resilience assessments to conserve and protect coastal ecosystems and to help build resilience in vulnerable ecosystems facing multiple threats.

The aim of this study is to investigate the effects of key environmental drivers (temperature, light, nutrient availability and grazing) that vary spatially and are expected to shift because of global change, on the resilience of foundational coastal ecosystems. Seagrass meadows, among the most threatened ecosystems worldwide^33^, were used as a model system. Since seagrass species traits, as well as the timing and temporal and spatial scale of the disturbance play a large role in determining resilience^34,35^, we standardized these factors within a regionally coordinated experiment. Resilience was assessed by measuring above- and belowground rates of seagrass recovery after a small-scale disturbance (complete biomass removal) across 10 sites in the Western North Atlantic, spanning >20° of latitude. We focused on the foundational species turtlegrass (*Thalassia testudinum*), as its range extends over a large region in the Western North Atlantic and varying recovery times have been reported across studies that varied in site characteristics and their methodology^36,37^. We tested the effects of nutrient fertilization (mimicking chronic eutrophication) on seagrass recovery at each site, and also measured important environmental covariates (temperature, light, grazing pressure – both fish and turtle). Generalized linear mixed models were then used to assess the separate and interactive effects of fertilization and environmental covariates on ecological resilience.

## Results

### Above- and belowground seagrass recovery

Aboveground recovery rates (calculated from biomass cores taken after one year of recovery) in unfertilized plots ranged 100-fold from 0.003 ± 0.001 (Bermuda) to 0.30 ± 0.06 (Bonaire) g DW m^-2^d^-1^ with an overall average of 0.06 ± 0.01 g DW m^-2^d^-1^ (Fig. 1A). Average belowground recovery rates per site ranged 10-fold from 0.04 ± 0.02 (Galveston, USA) to 0.42 ± 0.07 (Bonaire) g DW m^-2^d^-1^, with an overall average of 0.19 ± 0.03 g DW m^-2^d^-1^ (Fig. 1B).

**Figure 1.**
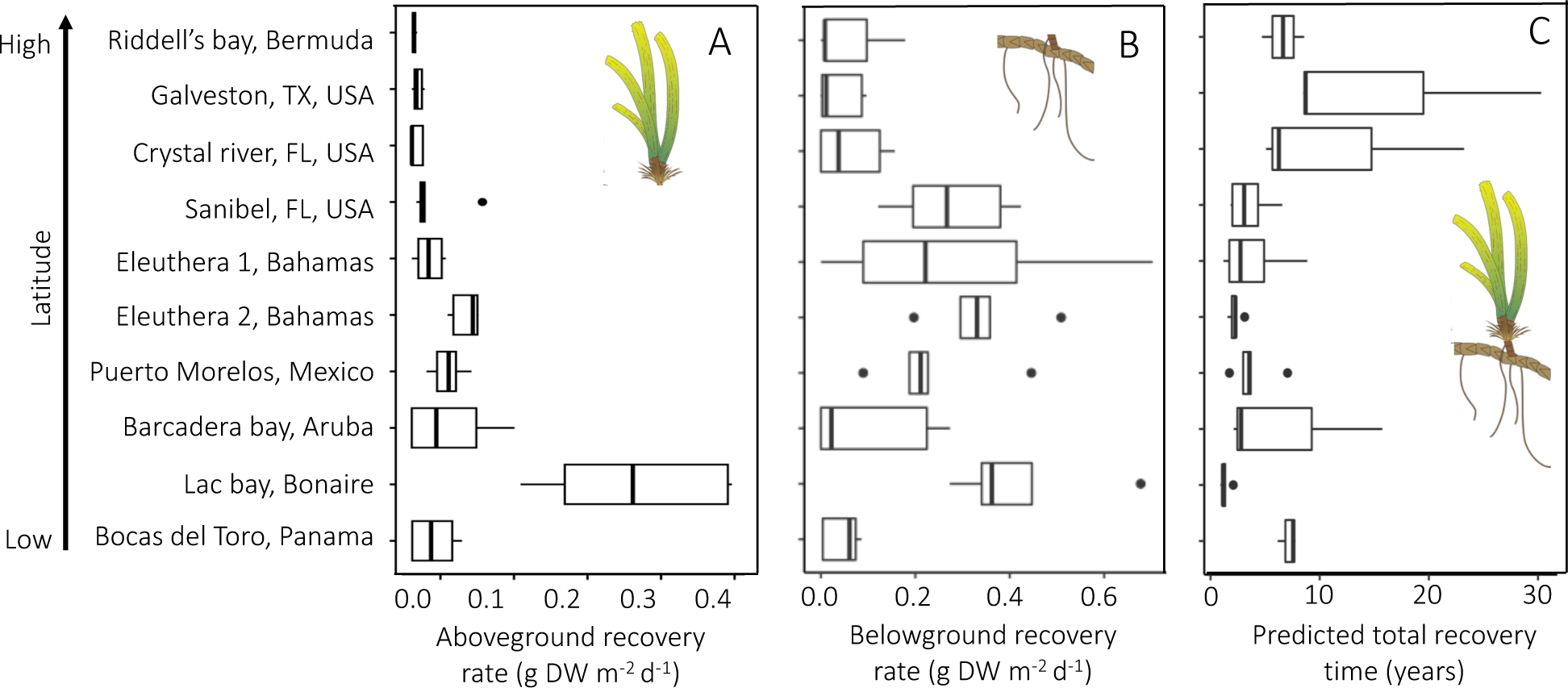
Boxplots of (a) aboveground and (b) belowground biomass recovery rates and (c) estimated total recovery time of turtlegrass in unfertilized plots in years. The order of the study sites corresponds to the latitudes from low latitude (bottom) to high latitude (top). Middle vertical lines of the boxes represent boxplot medians, left and right vertical lines represent the 25^th^ and 75^th^ percentiles, whiskers represent the smallest and largest measured values within the 1.5 interquartile range from the box and dots represent the outliers outside the interquartile range.

The percentage of above- and belowground biomass recovered in the unfertilized plots after one year was lowest for Crystal River, USA with 2.11 % ± 1.2 and 3.61 ± 1.9, respectively, and highest for Bonaire with 195.88 % ± 81.8 and 49.25 ± 6.5, respectively. Comparing above to belowground recovery rates per site and then averaging across sites, we found that aboveground biomass recovers to initial conditions 1.4 times faster than belowground biomass.

Additionally, we extrapolated the years needed to achieve full recovery (restoring the values to those of initial measurements), with a slightly reduced dataset due to removal of the plots that had zero recovery. Years needed for full recovery (both above- and belowground) were lowest for Bonaire (1.5 ± 0.18 years) and highest for Galveston (13.7 ± 3.16 years), with an average of 4.3 years across all sites.

The percentage of shoots and aboveground biomass recovered after one year were positively correlated with the percentage of belowground biomass recovered after one year (Pearson’s correlation test, R^2^ = 0.38, p < 0.001 and R^2^ = 0.40, p = 0.001 respectively).

There was no difference in seagrass cover in the control plots between the start and end of the experiment for all sites (p > 0.05), except Galveston which, because of high turbidity, could not be included (see Methods).

### The effects of fertilization and environmental factors on seagrass shoot recovery

There was no significant effect of fertilization on the percentage of recovered shoots of turtlegrass after one year (Table 1). Across the geographic range of turtlegrass, the percentage of shoots recovered increased with temperature (p = 0.03, Fig. 2). Significant interactions were observed between fertilization and both types of grazing pressure (fish and turtles). The percentage of recovered shoots increased with increasing fish grazing pressure in the unfertilized, but not in the fertilized plots (p = 0.002). With increasing turtle grazing pressure in the fertilized plots, the percentage of recovered shoots decreased, while no relationship was found in the unfertilized plots (p = 0.013). Fertilization therefore reduced shoot recovery at high grazing pressure for both types of grazers (see Supplementary Fig. 1. For trends in fish and turtle herbivory across sites).

**Table 1.**
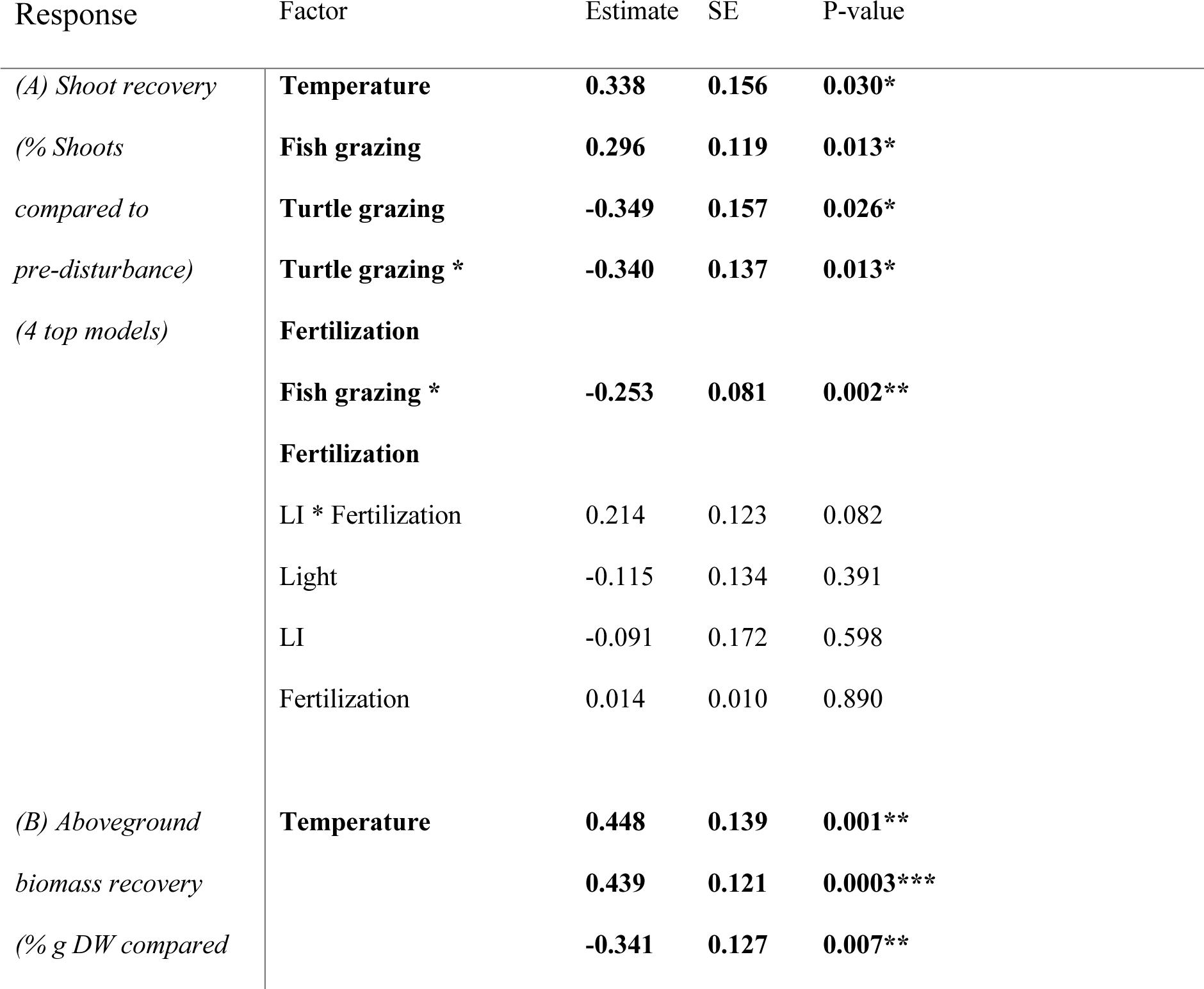
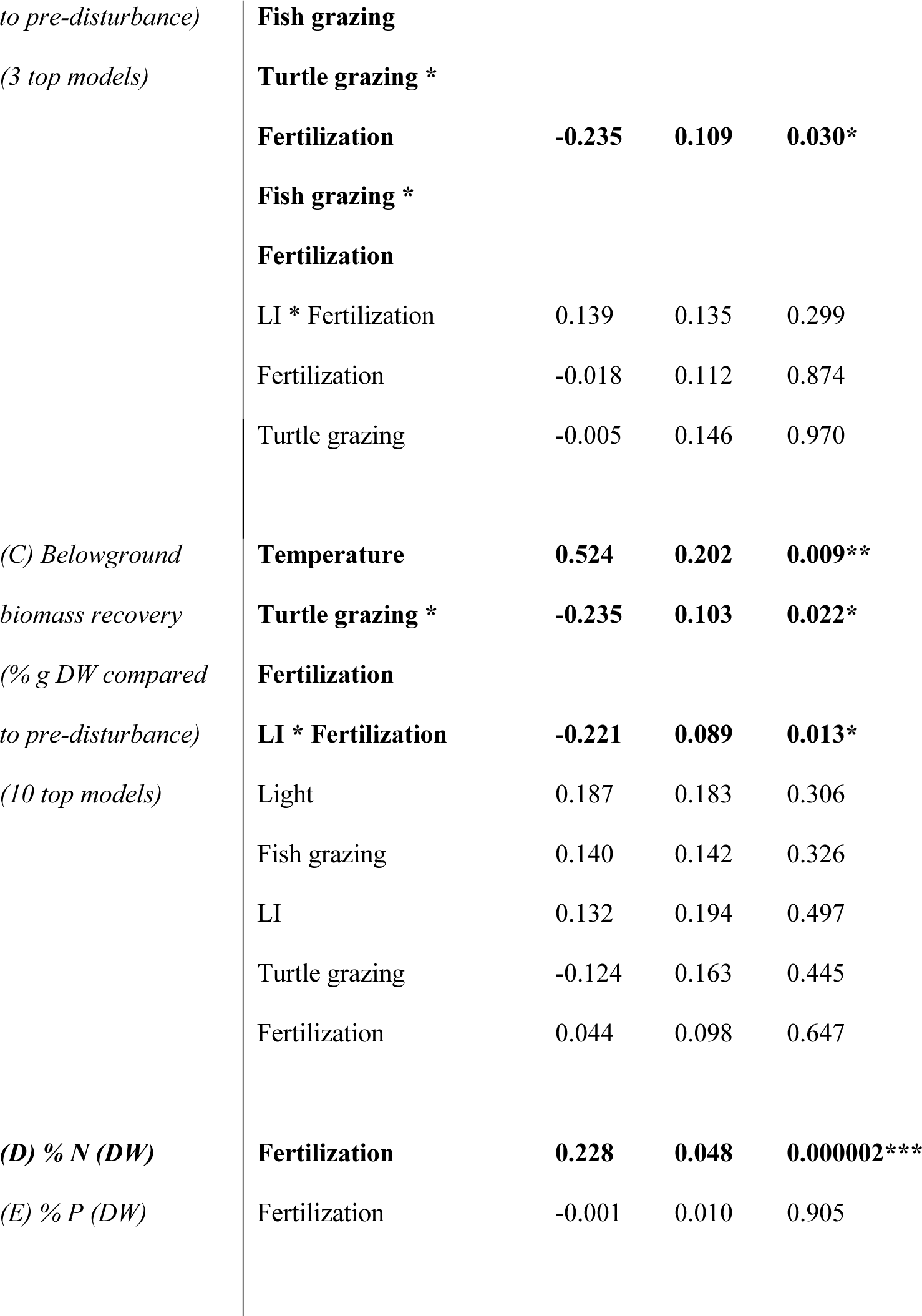
Statistical results for averaged generalized linear mixed models testing the impact of fertilization treatments and environmental drivers on seagrass recovery and nutrient content. The number of top models (≤ Δ2 AICc) is reported, along with the coefficient estimates and standard errors of the standardized regressors. Temperature is the average yearly water temperature at canopy level. Turtle and fish grazing is a grazing index assessed from the leaves. LI is the nutrient limitation index. Light is the yearly average input of light in the system. Nutrient fertilization was simulated by adding both N and P to the water column. Since only one factor, fertilization, was tested against nitrogen and phosphorus content, model averaging was not performed on those two models. Significance codes: ***p<0.0001, **p<0.01, *p<0.05.

**Figure 2.**
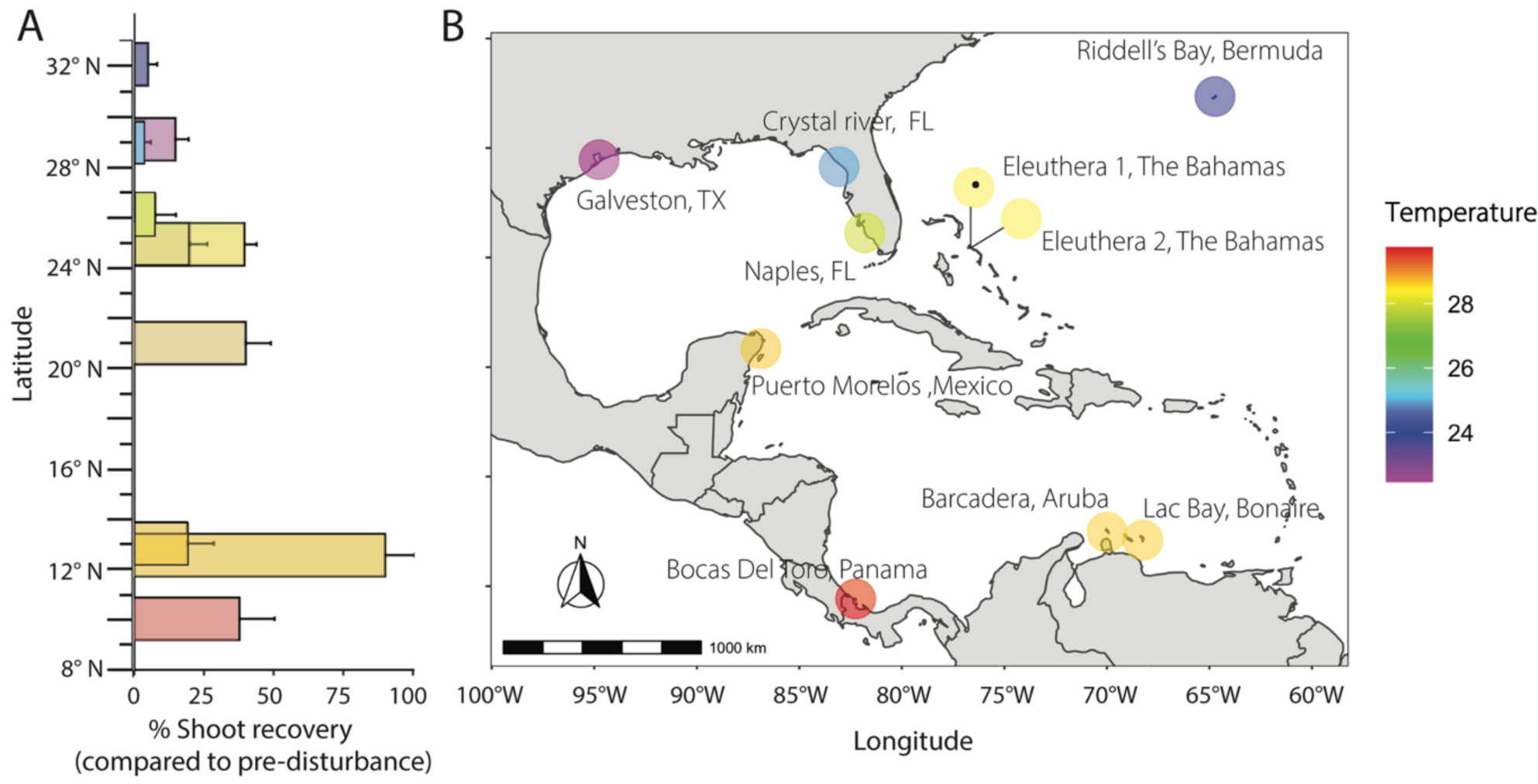
(A) Field measurements of seagrass shoot recovery (% compared to pre-disturbance ± SE) in unfertilized plots along a latitudinal gradient, with (B) a map of our study sites. Average annual water temperatures are visualized in color on the bar chart and on the map.

Shoot recovery rate decreased with seasonality, but not with latitude (Supplementary Table 1, 2).

### The effects of fertilization and environmental drivers on seagrass above- and belowground biomass recovery

Similar to shoot recovery, the percentage of aboveground biomass of turtlegrass recovered one year after disturbance increased with temperature (p = 0.001) (Fig. 3, Table 1), and significant interactions were found between fertilization and grazing pressure. The positive relationship between fish grazing pressure and aboveground biomass recovery was significantly reduced by fertilization (p < 0.001). Additionally, the positive relationship between turtle grazing pressure and aboveground biomass recovery was significantly reduced by fertilization, resulting in a negative relationship (p = 0.007). Aboveground biomass recovery decreased with latitude and with seasonality (Supplementary Table 1, 2).

**Figure 3.**
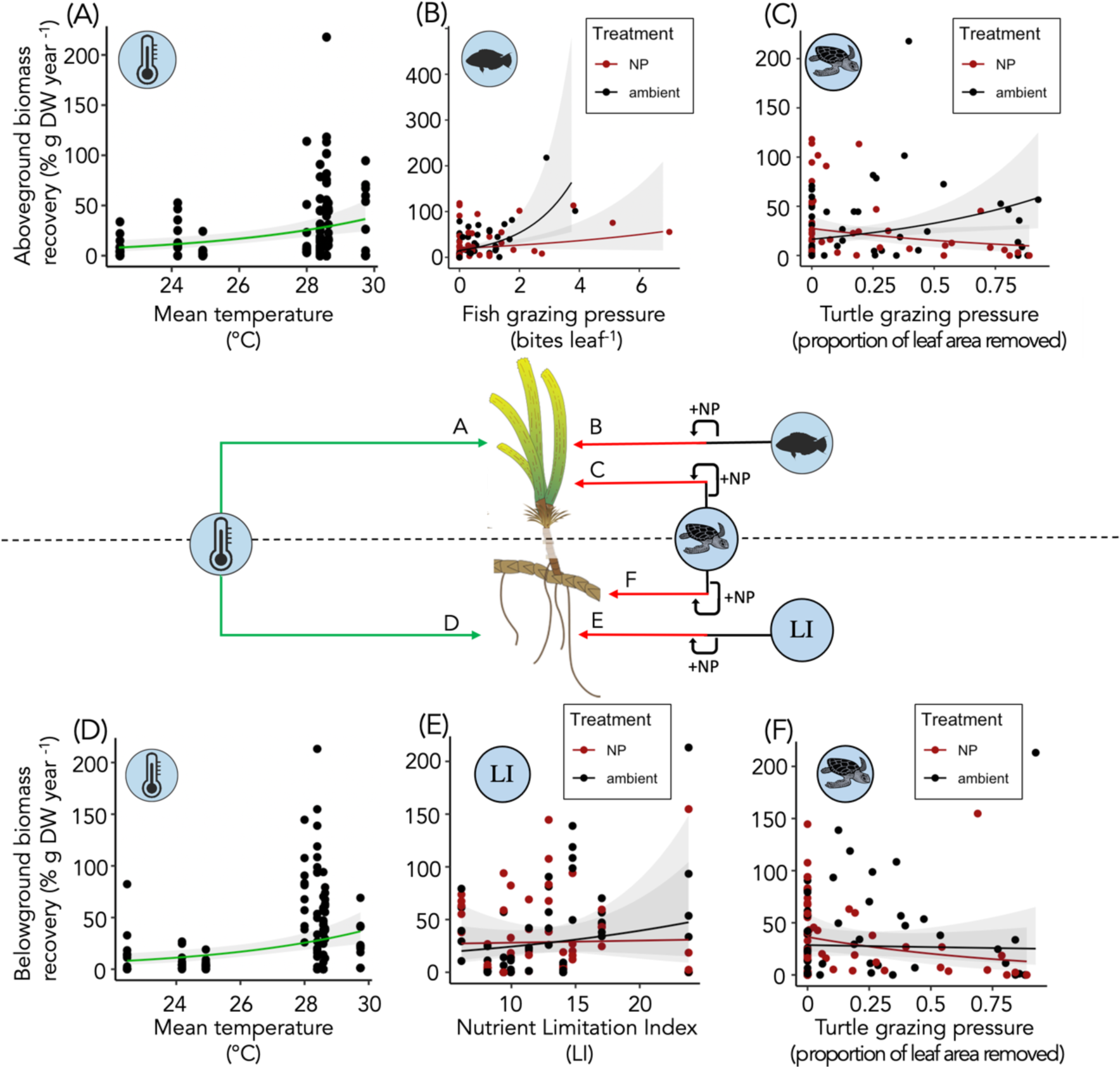
Summary of the results of the averaged generalized linear mixed models for above and belowground biomass recovery (% compared to pre-disturbance). The line represents the average value of the model response, with the 95% confidence interval, and is plotted on top of the measured data points. Arrows point from an environmental factor to either above or belowground biomass recovery and indicate a positive (green) or negative (red) significant or neutral (black) impact on recovery rates either as main effect or in the interaction with fertilization (+ NP) based on the coefficients from the models where p-values < 0.05. For example, fish grazing pressure in nutrient enriched plots decreased aboveground biomass recovery relative to ambient conditions. Arrow letters correspond to the plots (A-F). Averaged model results are presented in Table 1.

The percentage of belowground biomass recovered one year after disturbance also increased with temperature (p = 0.009) (Fig. 3, Table 1). Additionally, significant interactions were found between the nutrient limitation index (LI) and the fertilization treatment (p = 0.013) and between turtle grazing and the fertilization treatment (p = 0.022), indicating that fertilization decreased belowground biomass recovery when higher levels of nutrient limitation or turtle grazing were present. Belowground biomass recovery decreased with seasonality, but not with latitude (Supplementary Table 1, 2).

Fertilization increased leaf N content (p < 0.001), but not leaf P content (p = 0.91) (Table 1) in seagrass leaves taken from biomass cores at the end of the experiment (see Supplementary Fig. 2 for trends in %N, %P, C:N and C:P across sites).

When we compared the response of static versus dynamic indicators to the environmental drivers, we found that temperature would not have turned up as an important factor had we focused on static indicators (Supplementary Table 3). None of the drivers had a significant impact on static shoot density.

## Discussion

The capacity of coastal ecosystems to recover after disturbances depends on various local environmental factors, most of which are increasingly affected by global change^38,39^. Here, we evaluated the ecological resilience of the widespread (sub)tropical marine plant turtlegrass (*Thalassia testudinum*) by measuring the response to disturbance across its geographic range and along gradients in environmental factors. Our results provide the first experimental evidence that both above- and belowground recovery rates of a marine plant increase with temperature after a disturbance. Specifically, we found that cooler temperatures at subtropical sites may limit resilience, increasing vulnerability to disturbances and potential seagrass meadow collapse^23,24^. Fish and turtle herbivory influenced aboveground recovery, depending on local nutrient availability. The potential for increased resilience due to increasing temperatures and reported interactions between grazers and local nutrient input could help tropical coastal foundation species at the cooler edges of their range to resist change and recover from future disturbances anticipated under global change scenarios.

By measuring dynamic indicators (i.e., recovery rates) instead of traditional static indicators (i.e., shoot density, percent cover) we found that a combination of temperature and the interaction between herbivory and fertilization drives the resilience of turtlegrass. Temperature increased seagrass shoot, aboveground and belowground biomass recovery but did not affect static aboveground biomass, and none of the measured environmental drivers significantly impacted static shoot density, which, along with cover, is commonly used as a seagrass response indicator. These results indicate that dynamic indicators are more responsive to environmental factors and better predict the responses of seagrasses to future disturbance. Additionally, we found that aboveground biomass recovered 1.4 times faster than belowground biomass, indicating that aboveground biomass production is likely followed by belowground biomass recovery. We want to highlight that belowground biomass recovery responded differently to climate-change related drivers compared to shoot and aboveground biomass recovery. Therefore, dynamic responses of aboveground variables may not be representative of the whole plant, and we recommend incorporating dynamic measures of belowground parts to understand the health and resilience of coastal foundation species^40^ and their adaptability to a changing environment^22^.

The positive relationship found between temperature and seagrass recovery contrasts with the negative effects of prolonged high temperatures that are regularly reported for seagrasses^41,42^, and other coastal foundation species^38,43^. While temperate seagrass meadows are especially vulnerable to warming^10^, turtlegrass, primarily found at tropical latitudes, may benefit from warming at the edges of its habitat range. Previous studies suggest mild temperature increases enhance seagrass photosynthetic rate^44^ and shoot formation through clonal growth^45^. However, beyond certain temperature thresholds, respiration can exceed photosynthesis, reducing growth rates^46^ and potentially causing collapse for species growing near the upper limits of their thermal distribution or in areas prone to heatwaves^10,47^. Our study suggests that within the examined temperature range, which did not include heatwaves, slight increases in mean annual temperature can enhance the recovery potential of subtropical seagrasses, just as has been found for salt marsh plants in a warming experiment^48^. Experimental testing of increasing temperature impacts on subtropical seagrass resilience is needed to confirm the correlative relationships found in this study and to assess whether within-site variations in temperature can also drive recovery rates.

Subtropical sites experience higher seasonality, due to low surface irradiance and temperatures in winter, and therefore shorter growing seasons^49^. We found that higher annual temperature variability, similar to lower annual temperature, reduces recovery rates. A longer growing season due to global warming may therefore increase seagrass resilience at subtropical latitudes, depending on local grazing pressure and light and nutrient availability. Recent work has demonstrated that subtropical seagrasses in the Western North Atlantic are increasingly sensitive to high grazing pressure and that light can play a key role in regulating seagrass response to overgrazing^16^. However, in this study, we found instead that temperature played a stronger role. Several reasons may account for this distinction across studies, such as the use of slightly varying metrics (leaf production vs whole shoot recovery). However, in sum, both studies document an increased vulnerability of a tropical seagrass at its cooler range boundary.

Our results highlight the importance of distinguishing types of grazers and their impact on seagrass recovery rates, corresponding to findings from a single-site experiment^50^. Grazing pressure is expected to rise, especially in the subtropics, due to the indirect effects of rising temperatures on herbivore habitat range and metabolism^15,51^. We found that fish grazing can positively impact aboveground recovery, likely due to compensatory growth or algae control^52^. An alternative explanation is that, similar to terrestrial grasslands, grazing may open up the canopy thereby increasing light availability for growth of shorter vegetation^53^. Migrating fishes from tropical to subtropical sites may therefore increase local meadow resilience, up to a herbivore density limit where intensive grazing prevents regrowth^54^. For larger herbivores such as green turtles, we found that grazing reduced the recovery in fertilized plots. At one of our sites on Eleuthera, heavy turtle grazing^55^ resulted in a > 50 % reduction in shoot recovery compared to a nearby ungrazed site. Our results, combined with reports of increasing overgrazing events at subtropical sites in the Western Atlantic^56,57^ suggest reduced meadow resilience and ecosystem functioning in turtle-dense environments^58,59^.

Theory suggests that the capacity of plant communities to recover after disturbances likely depends on local nutrient status^60,61^. In our study, nutrient limitation positively affected belowground biomass recovery in unfertilized plots, but this effect was significantly reduced by fertilization, potentially because seagrass invests less in belowground recovery when nutrients are abundant^16,62^. Additionally, grazing by both turtles and fish reduced the aboveground recovery in fertilized plots, likely due to increased grazing pressure on nutrient-enriched leaves^63^. Fertilization-induced grazing pressure by turtles also decreased belowground recovery rates. Thus, high latitude sites, currently less resilient due to temperature and seasonality effects, may become more vulnerable to consumer pressure fueled by eutrophication, as was found for other coastal wetlands^64^. Therefore, it is important to monitor how subtropical seagrasses respond to expected increases in temperature and grazing pressures as well as to assess their carbohydrate reserves and (seasonal) light availability to determine if they will be able to maintain resilience under high grazing pressure^16^

Our study demonstrates that 1) temperature can increase (sub) tropical seagrass resilience, 2) nutrient fertilization combined with fish or turtle herbivory can decrease seagrass meadow resilience, and 3) measuring belowground biomass in addition to aboveground biomass is essential for understanding the resilience of coastal foundation species. Dynamic indicators like recovery rate are essential for estimating ecosystem resilience, and our findings underscore the value of replicated experiments across large environmental gradients. Ecologically based strategies are needed to enhance the resilience of coastal ecosystems and maintain their roles as ecosystem engineers in a changing world.

## Methods

### Study site

This study was part of a larger coordinated research program, the *Thalassia* Experimental Network (TEN), which consisted of a series of sites across the geographic range of turtlegrass (*Thalassia testudinum*) in the Western North Atlantic (9-32 °N)^16^. At each site, seagrass meadows were selected based on the following criteria: (1) depth of < 4 m, (2) dominated by turtlegrass (> 50 % relative abundance), and (3) a minimum area of 25 m x 25 m. Due to logistics, this experiment was performed at 9 out of the 13 sites that were part of TEN (Supplementary Table 4, Fig. 2). To improve the latitudinal balance of the set-up we added one site that was not in the original network: Barcadera Bay, Aruba. Additionally, the original TEN site on Eleuthera became heavily grazed by turtles due to the presence of experimental cages and was therefore not representative of the surrounding seagrass seascape^55^. We established a second additional site (Eleuthera 2) outside of the grazing patch.

### Experimental design

Ten experimental plots (0.25 x 0.25 m, at least 2 m apart in a randomized design) were established at each site in the fall of 2018 (Sept – Nov, Supplementary Table 4). In each plot, a disturbance was created by removing all above – and belowground biomass within a 15 cm diameter circle, 20 cm deep. After the biomass core was collected, the void was filled with local sediment, and bamboo skewers (∼6 per plot) were used to mark the exact border where the biomass core had been collected. The species turtlegrass mainly recovers through clonal growth via elongation of horizontal rhizomes^65^. A replicated experiment was conducted at each site with two treatments, nutrient fertilized and unfertilized conditions (N = 5 plots per treatment). Every two weeks to two months (depending on logistics), the number of shoots regrown in the void was counted, to investigate whether the shoot establishment rate was linear throughout the year. After about one year (10-14 months after disturbance) all biomass that had recovered within the marked void was collected.

We set up paired control plots by assessing the seagrass cover in the 0.06 m^2^ area directly surrounding unfertilized plots (N = 3). In these control plots, we assessed seagrass cover from photos taken both at the initiation of the experimental perturbation, and at the end of the experiment. Due to high turbidity, we were unable to include Galveston in this analysis, but personal observations confirm no major changes at this site in seagrass cover for the duration of the experiment (pers. obs. JAG, ARA)

Both at the start and end of the experiment, all seagrass material was stored in a cooler and processed within 24 hours. The shoots were separated from the belowground biomass, leaves were scraped clean of epiphytes, and the above and belowground material were dried separately in an oven at 60 °C. The number of shoots within the biomass core was recorded as well as the dry weight of the above- and belowground biomass per plot.

Fertilization treatments were established by attaching a fiberglass mesh bag containing 300 g of slow-release Osmocote fertilizer (Everris NPK 14:14:14) 30 cm above the sediment to a pole, at a corner of each plot, following^16^. Bags were replaced monthly to ensure consistent enrichment.

### Environmental drivers of seagrass recovery

We measured several environmental factors at the site level as candidate drivers for seagrass recovery. Underwater loggers deployed in the seagrass canopy (HOBO UA-002064) recorded the water temperature every 6 minutes at each site. From these measurements an average annual temperature was calculated, as well as the seasonality (SD of temperature among months). Light intensity was measured by a light sensor (Odyssey Submersible PAR Logger) deployed at the same location, with the same measuring interval and duration as the temperature loggers and averaged annually. Since sites may be either P- or N-limited^66^, we used an index to indicate the overall magnitude of nutrient limitation. The Limitation Index (LI) was calculated as the absolute deviation of leaf molar N:P from the balanced 30:1 ratio^16^. LI indicates ambient nutrient availability, where higher LI values signal a larger degree of either N- or P-limitation. Ambient leaf N and P content was obtained by analyzing the green leaf tissue from the unfertilized (N = 5) plots at the start of the experiment. Additionally, all green leaf material of both unfertilized (N = 5) and fertilized plots (N = 5) obtained at the end of the experiment was analyzed for nutrients to assess the impact of nutrient enrichment on leaf N and P content. Dried leaf material was homogenized to a fine powder using a mortar and pestle. The leaf material was subsequently analyzed for nitrogen content on an elemental analyzer (Thermo Flash 1112), and for phosphorus content on an autoanalyzer (SKALAR San++) after a digestion using sulphuric acid and selenium^67^.

To quantify herbivory pressure, we estimated both megaherbivore (turtle) and mesoherbivore (fish) grazing pressure per plot. Fish grazing on turtlegrass results in crescent shaped bitemarks from the sides and top of the leaves (Supplementary Fig. 3a). Therefore, fish grazing pressure was estimated by counting the average number of fish (crescent) grazing marks per shoot of (a maximum of) 10 shoots collected in each plot at the end of the experiment (fall 2019). Turtles crop the leaves from above resulting in a straight cut (Supplementary Fig. 3b). Therefore, turtle grazing pressure was estimated by calculating the proportion of leaf area that was removed in each of the plots relative to the mean leaf area of the unfertilized caged plots of the TEN experiment^16^. The outcome was validated by comparing it with known turtle abundances at the study sites (Pers. obs. LMRB, AMM, FOHS, SAM). Estimates were based on seagrass leaf grazing marks instead of known fish or turtle densities in the area because these more accurately represent local grazing impact as within a given meadow there can be local heterogeneity in grazing pressure^55,63^.

### Data analysis

We used multi-model inference to examine which local and across-site environmental factors were important for our recovery response variables which were based on measured shoot abundance, aboveground biomass and belowground biomass. Because the timing of the end-harvest varied across sites (between 293 – 433 days after the start of disturbance), the number of shoots, aboveground biomass and belowground biomass recovered at the end of the experiment were standardized to 365 days. For this, we assumed linear growth, which was confirmed using regression analysis of the shoot abundance data over time after experimental disturbance (Supplementary Table 5) and by a recent review stating most seagrass recovery studies provide evidence of linear recovery over time^27^. The percentage recovered was calculated by dividing the plot-specific response variables after 1 year by the pre-disturbance values measured, and multiplying this by 100. Years needed until full recovery was calculated by dividing the start total biomass by the end total biomass, multiplied by the duration of each experiment, and divided by 365 days.

Latitude and seasonality were both correlated with average annual temperature and therefore excluded from the main models (Supplementary Table 6), and are presented in the supporting information (Supplementary Table 1, 2). Average temperature was selected over latitude and seasonality since it is the candidate driver reported to increase with global warming.

For all response variables (shoot recovery, aboveground and belowground recovery), we included the covariates ‘temperature’, ‘fish grazing’, ‘turtle grazing’, ‘light’ and ‘LI’. The models also included fertilization as a fixed factor to observe any significant interactions between fertilization and fish herbivory, turtle herbivory, and LI (Supplementary Equation 1). We standardized our covariate values by subtracting the mean and dividing by the standard deviation. All covariates had variance inflation factors <5, indicating low collinearity. We fitted the full models for all response variables using generalized linear mixed models (GLMMs) with site as a random effect and a Tweedie distribution used for continuous data with non-normal distributions and zero inflation (our response variables had between 13 – 26% zeroes and were tested for zero-inflation using the DHARMa package) using the glmmTMB package. All full models were examined for model fit by plotting the residuals versus the fitted values, the fitted values versus the observed data and the residuals versus the treatment ‘fertilization’. The model fit, specifically the ability of the models to cope with the large numbers of zeroes, as well as outliers, dispersion and uniformity were tested using the DHARMa package. We ranked the resulting potential models with AICc using the ‘dredge’ function in the MuMIn package in R. Because the top models were performing equally well, we performed model averaging to arrive at consistent parameter estimates of the most important explanatory variables in the full GLMM, by averaging a set of top models which share similarly high levels of parsimony (see Supplementary Table 7 for the selection of top models). We defined the top models as those that fell within 2 AIC units of the model with the lowest AIC value, as is recommended when factors may have weak interactions with the response^68^ with the model.avg function in the MuMIn package, and we present the conditional averages. Standardized coefficient plots are visualized in Supplementary Fig. 4. For data visualization of above- and belowground biomass recovery, we created a dataset using the ‘predict’ function for each specific significant variable while the remaining variables were set at their average value.

To test whether fertilization increased leaf N and P content, we fitted a linear mixed effects model with a gaussian distribution (using glmmTMB) to plot-specific leaf N and P data, with site as a random effect and fertilization as a fixed factor. Model validation was conducted as described above.

To test the difference in effect of the environmental factors on traditional static indicators versus dynamic indicators, we compared the response of static indicators: aboveground biomass (g DW m^-2^) and shoot density (shoots m^-2^) as measured before the experimental disturbance, to dynamic indicators: aboveground biomass recovery and shoot recovery percentages as obtained at the end of the experiment in the unfertilized plots (total of 50 plots). For shoot density and aboveground biomass linear mixed models were used (using lme4 package) and for shoot and aboveground biomass recovery generalized linear mixed models with a Tweedie distribution (using glmmTMB).

To investigate the relationships among the above and belowground seagrass recovery response variables, we performed correlation analysis using the ‘cor’ function in R.

To test whether the seagrass cover at the site remained stable over the course of the experiment we compared the paired control plots between the start and end of the experiment using pairwise t-tests with a Bonferroni correction. All data analyses were performed in R (v.4.2.2).

## Supporting information

Supplemental Figs. 1-4, Supplemental Tables 1-7 and Supplemental Equation 1

## Acknowledgments

We thank the many staff, students and volunteers who contributed to the field and laboratory research of this study. We thank Scott Jones, Zachary Foltz, Skylar Carlson, Iris Segura-Garcia, Maggie Johnson, Audrey Looby, Olivia Carmack and David Branson at the Smithsonian Marine Station; Kathryn Coates at the Bermuda site; Sabine Engel and Jessica Johnson at the Bonaire site; Scott Alford, Theresa Gruninger, Audrey Looby, Cayla Sullivan, Sawyer Downey, Whitney Scheffel, Jamila Roth and Tim Jones at the Crystal River site; Tom Gluckman, Cameron Ragusa, Isabella Primrose Hartman, William F. Bigelow and Matheo Albury at the Eleuthera site, Ashley E. McDonald at the Galveston site, and M. Guadelupe Barba Santos at the Puerto Morelos site. This work was conducted under the following permits: permit #s MAMR/FIS/17 and MAMR/FIS/9 at Eleuthera; permit #558/2015-201500762 at Bonaire; permit #s SE/AP-23-17 and SE/AO-1-19 at Bocas del Toro. Funding for this project was provided by the US National Science Foundation (OCE-1737247 to JEC, AHA, and VJP, OCE-2019022 to JEC, OCE-1737144 to KLH, and OCE-1737116 to JGD). MJAC was supported by NWO-Veni grant 181.002. FOHS was supported by the 2019 Ecology Fund of the Royal Netherlands Academy of Arts and Sciences and by the Netherlands Earth System Science Centre (NESSC). This is contribution #X from the Coastlines and Oceans Division of the Institute of Environment at Florida International University.

## References

1. Folke, C. et al. Regime shifts, resilience, and biodiversity in ecosystem management. Annu. Rev. Ecol. Evol. Syst 35, (2004).

2. Dirzo, R. et al. Defaunation in the Anthropocene. Science (1979) 345, 401–406 (2014).

3. Woodruff, J. D., Irish, J. L. & Camargo, S. J. Coastal flooding by tropical cyclones and sea-level rise. Nature 2013 504:7478 504, 44–52 (2013).

4. IPCC. Climate Change 2022: Impacts, Adaptation and Vulnerability. https://www.ipcc.ch/report/sixth-assessment-report-working-group-ii/ (2022).

5. He, Q. & Silliman, B. R. Climate change, human impacts, and coastal ecosystems in the Anthropocene. Current Biology 29, R1021–R1035 (2019).

6. Gissi, E. et al. A review of the combined effects of climate change and other local human stressors on the marine environment. Science of The Total Environment 755, 142564 (2021).

7. James, R. K. et al. Climate change mitigation by coral reefs and seagrass beds at risk: How global change compromises coastal ecosystem services. Science of The Total Environment 857, 159576 (2023).

8. Temmink, R. J. M. et al. Recovering wetland biogeomorphic feedbacks to restore the world’s biotic carbon hotspots. Science 376, 6593 (2022).

9. Spalding, M. D. et al. The role of ecosystems in coastal protection: Adapting to climate change and coastal hazards. Ocean Coast Manag 90, 50–57 (2014).

10. Marbà, N., Jordà, G., Bennett, S. & Duarte, C. M. Seagrass thermal limits and vulnerability to future warming. Front Mar Sci 9, (2022).

11. Saintilan, N., Wilson, N. C., Rogers, K., Rajkaran, A. & Krauss, K. W. Mangrove expansion and salt marsh decline at mangrove poleward limits. Glob Chang Biol 20, 147–157 (2014).

12. Seidl, R. et al. Forest disturbances under climate change. Nature Climate Change 7, 395–402 (2017).

13. Knutson, T. R. et al. Tropical cyclones and climate change. Nature Geoscience 3, 157–163 (2010).

14. Serrano, O. et al. Impact of marine heatwaves on seagrass ecosystems. 345–364 (2021).

15. Vergés, A. et al. Long-term empirical evidence of ocean warming leading to tropicalization of fish communities, increased herbivory, and loss of kelp. Proc Natl Acad Sci U S A 113, 13791–13796 (2016).

16. Campbell, J. E. et al. Herbivore effects increase with latitude across the extent of a foundational seagrass. Nature Ecology & Evolution 2024 8, 663–675 (2024).

17. Horta, P. A. et al. Marine eutrophication: overview from now to the future. in Anthropogenic Pollution of Aquatic Ecosystems 157–180 (2021).

18. Deegan, L. A. et al. Coastal eutrophication as a driver of salt marsh loss. Nature 490, 388– 392 (2012).

19. Van Nes, E. H. & Scheffer, M. Slow recovery from perturbations as a generic indicator of a nearby catastrophic shift. American Naturalist 169, 738–747 (2007).

20. Pazzaglia, J. et al. Does warming enhance the effects of eutrophication in the seagrass *Posidonia oceanica*? Front Mar Sci 7, 1067 (2020).

21. Holling, C. S. & Gunderson, L. H. Resilience and adaptive cycles. Panarchy: Understanding transformations in human and natural systems 25–62 (2002).

22. Cole, L. E. S., Bhagwat, S. A. & Willis, K. J. Recovery and resilience of tropical forests after disturbance. Nat Commun 5, (2014).

23. Scheffer, M., Carpenter, S. R., Dakos, V. & Van Nes, E. H. Generic indicators of ecological resilience: inferring the chance of a critical transition. Annu Rev Ecol Evol Syst 46, 145–167 (2015).

24. van de Leemput, I. A., Dakos, V., Scheffer, M. & van Nes, E. H. Slow recovery from local disturbances as an indicator for loss of ecosystem resilience. Ecosystems 21, 141–152 (2018).

25. Oliver, T., et al. Biodiversity and resilience of ecosystem functions. Trends in Ecology & Evolution 30, 673–684 (2015).

26. Castagno, K. A. et al. Resistance, resilience, and recovery of salt marshes in the Florida Panhandle following Hurricane Michael. Scientific Reports 2021 11:1 11, 1–10 (2021).

27. Tassone, S. J., Ewers Lewis, C. J., McGlathery, K. J. & Pace, M. L. Seagrass ecosystem recovery: Experimental removal and synthesis of disturbance studies. Limnol Oceanogr 69, 1593–1605 (2024).

28. Vonk, J. A., Christianen, M. J. A., Stapel, J. & O’Brien, K. R. What lies beneath: Why knowledge of belowground biomass dynamics is crucial to effective seagrass management. Ecol Indic 57, 259–267 (2015).

29. Nyman, J. A., Walters, R. J., Delaune, R. D. & Patrick, W. H. Marsh vertical accretion via vegetative growth. Estuar Coast Shelf Sci 69, 370–380 (2006).

30. Yang, H. & Li, T. Responses of above-and belowground carbon stocks to degraded and recovering wetlands in the Yellow river delta. Front Ecol Evol (2022).

31. Battisti, D. De & Griffin, J. Below-ground biomass of plants, with a key contribution of buried shoots, increases foredune resistance to wave swash. Ann Bot (2022).

32. Infantes, E. et al. Seagrass roots strongly reduce cliff erosion rates in sandy sediments. Mar Ecol Prog Ser 700, 1–12 (2022).

33. Dunic, J. C., Brown, C. J., Connolly, R. M., Turschwell, M. P. & Côté, I. M. Long-term declines and recovery of meadow area across the world’s seagrass bioregions. Glob Chang Biol 27, 4096–4109 (2021).

34. Sanmartí, N., M. Ricart, A., Ontoria, Y., Pérez, M. & Romero, J. Recovery of a fast-growing seagrass from small-scale mechanical disturbances: Effects of intensity, size and seasonal timing. Mar Pollut Bull 162, 111873 (2021).

35. O’Brien, K. R. et al. Seagrass ecosystem trajectory depends on the relative timescales of resistance, recovery and disturbance. Mar Pollut Bull 134, 166–176 (2018).

36. Hammerstrom, K. K., Kenworthy, W. J., Whitfield, P. E. & Merello, M. F. Response and recovery dynamics of seagrasses *Thalassia testudinum* and *Syringodium filiforme* and macroalgae in experimental motor vessel disturbances. Mar Ecol Prog Ser 345, 83–92 (2007).

37. Dawes, C. J., Andorfer, J., Rose, C., Uranowski, C. & Ehringer, N. Regrowth of the seagrass *Thalassia testudinum* into propeller scars. Aquat Bot 59, 139–155 (1997).

38. Wernberg, T. et al. Decreasing resilience of kelp beds along a latitudinal temperature gradient: potential implications for a warmer future. Ecol Lett 13, 685–694 (2010).

39. Jones, S. F. et al. Stress gradients interact with disturbance to reveal alternative states in salt marsh: Multivariate resilience at the landscape scale. Journal of Ecology 109, 3211–3223 (2021).

40. Lam, V. Y. Y., Doropoulos, C. & Mumby, P. J. The influence of resilience-based management on coral reef monitoring: A systematic review. PLoS One 12, e0172064 (2017).

41. Aoki, L. R. et al. Seagrass recovery following marine heat wave influences sediment carbon stocks. Front Mar Sci 7, 1170 (2021).

42. Strydom, S. et al. Too hot to handle: Unprecedented seagrass death driven by marine heatwave in a World Heritage Area. Glob Chang Biol 26, 3525–3538 (2020).

43. Smale, D. A. Impacts of ocean warming on kelp forest ecosystems. New Phytologist 225, 1447–1454 (2020).

44. Lee, K.-S., Park, S. R. & Kim, Y. K. Effects of irradiance, temperature, and nutrients on growth dynamics of seagrasses: A review. J Exp Mar Biol Ecol 350, 144–175 (2007).

45. Lee, K. & Dunton, K. H. Production and carbon reserve dynamics of the seagrass *Thalassia testudinum* in Corpus Christi Bay, Texas, USA. Mar Ecol Prog Ser 143, 201–210 (1996).

46. Nguyen, H. M., Ralph, P. J., Marín-Guirao, L., Pernice, M. & Procaccini, G. Seagrasses in an era of ocean warming: a review. Biological Reviews 96, 2009–2030 (2021).

47. Wiens, J. J. Climate-related local extinctions are already widespread among plant and animal species. PLoS Biol 14, e2001104 (2016).

48. Smith, A. J., Noyce, G. L., Megonigal, J. P., Guntenspergen, G. R. & Kirwan, M. L. Temperature optimum for marsh resilience and carbon accumulation revealed in a whole-ecosystem warming experiment. Glob Chang Biol (2022).

49. Tussenbroek, B. I. van et al. Caribbean-wide, long-term study of seagrass beds reveals local variations, shifts in community structure and occasional collapse. PLoS One 9, e90600 (2014).

50. O’Dea, C. M., Lavery, P. S., Webster, C. L. & McMahon, K. M. Increased extent of waterfowl grazing lengthens the recovery time of a colonizing seagrass (*Halophila ovalis)* with implications for seagrass resilience. Front Plant Sci 13, 947109–947109 (2022).

51. Zarco-Perello, S. et al. Range-extending tropical herbivores increase diversity, intensity and extent of herbivory functions in temperate marine ecosystems. Funct Ecol 34, 2411–2421 (2020).

52. Valentine, J. F. & Heck, K. L. Herbivory in seagrass meadows: an evolving paradigm. Estuaries and Coasts 44, 491–505 (2021).

53. Borer, E. T. et al. Herbivores and nutrients control grassland plant diversity via light limitation. Nature 2014 508:7497 508, 517–520 (2014).

54. Bennett, S., Wernberg, T., Harvey, E. S., Santana-Garcon, J. & Saunders, B. J. Tropical herbivores provide resilience to a climate-mediated phase shift on temperate reefs. Ecol Lett 18, 714–723 (2015).

55. Smulders, F. O. H. et al. Green turtles shape the seascape through grazing patch formation around habitat features: Experimental evidence. Ecology 104, e3902 (2023).

56. Fourqurean, J. W., Manuel, S. A., Coates, K. A., Massey, S. C. & Kenworthy, W. J. Decadal monitoring in Bermuda shows a widespread loss of seagrasses attributable to overgrazing by the green sea turtle *Chelonia mydas*. Estuaries and Coasts 42, 1524–1540 (2019).

57. Rodriguez, A. R. & Heck, K. L. Approaching a tipping point? Herbivore carrying capacity estimates in a rapidly changing, seagrass-dominated Florida Bay. Estuaries and Coasts 44, 522–534 (2021).

58. Gangal, M. et al. Sequential overgrazing by green turtles causes archipelago-wide functional extinctions of seagrass meadows. Biol Conserv 260, 109195 (2021).

59. Christianen, M. J. A. et al. Seagrass ecosystem multifunctionality under the rise of a flagship marine megaherbivore. Glob Chang Biol 29, 215–230 (2023).

60. Boada, J. et al. Immanent conditions determine imminent collapses: nutrient regimes define the resilience of macroalgal communities. Proceedings of the Royal Society B: Biological Sciences 284, (2017).

61. Wasson, K. et al. Eutrophication decreases salt marsh resilience through proliferation of algal mats. Biol Conserv 212, 1–11 (2017).

62. Romero, J., Lee, K. S., Pérez, M., Mateo, M. A. & Alcoverro, T. Nutrient dynamics in seagrass ecosystems. in Seagrasses: Biology, Ecology and Conservation 227–254 (2006).

63. Smulders, F. O. H. et al. Fish grazing enhanced by nutrient enrichment may limit invasive seagrass expansion. Aquat Bot 176, 103464 (2022).

64. He, Q. & Silliman, B. R. Biogeographic consequences of nutrient enrichment for plant– herbivore interactions in coastal wetlands. Ecol Lett 18, 462–471 (2015).

65. van Tussenbroek, B. I. et al. The biology of Thalassia: Paradigms and recent advances in research. in Seagrasses: Biology, Ecology and Conservation 409–439 (Springer Netherlands, Dordrecht, 2006).

66. Fourqurean, J. W. et al. Seagrass abundance predicts surficial soil organic carbon stocks across the range of *Thalassia testudinum* in the Western North Atlantic. Estuaries and Coasts 46, 1280–1301 (2023).

67. Novozamsky, I., Houba, V. J. G., van Eck, R. & van Vark, W. A novel digestion technique for multi-element plant analysis. Commun Soil Sci Plant Anal 14, 239–248 (1983).

68. Grueber, C. E., Nakagawa, S., Laws, R. J. & Jamieson, I. G. Multimodel inference in ecology and evolution: Challenges and solutions. Journal of Evolutionary Biology vol. 24 699–711 (2011).

