## Supplemental Figs. 1-4, Supplemental Tables 1-7 and Supplemental Equation 1 for "Temperature drives seagrass recovery across the Western North Atlantic"

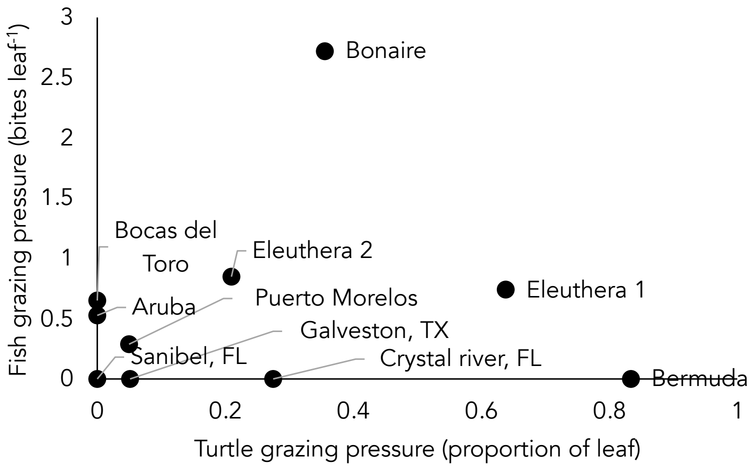


**Supplementary Figure S1.** Herbivory grazing pressure across sites. Site means for fish grazing pressure (number of bites per leaf) and turtle grazing pressure (proportion of leaf removed) as measured in unfertilized plots.

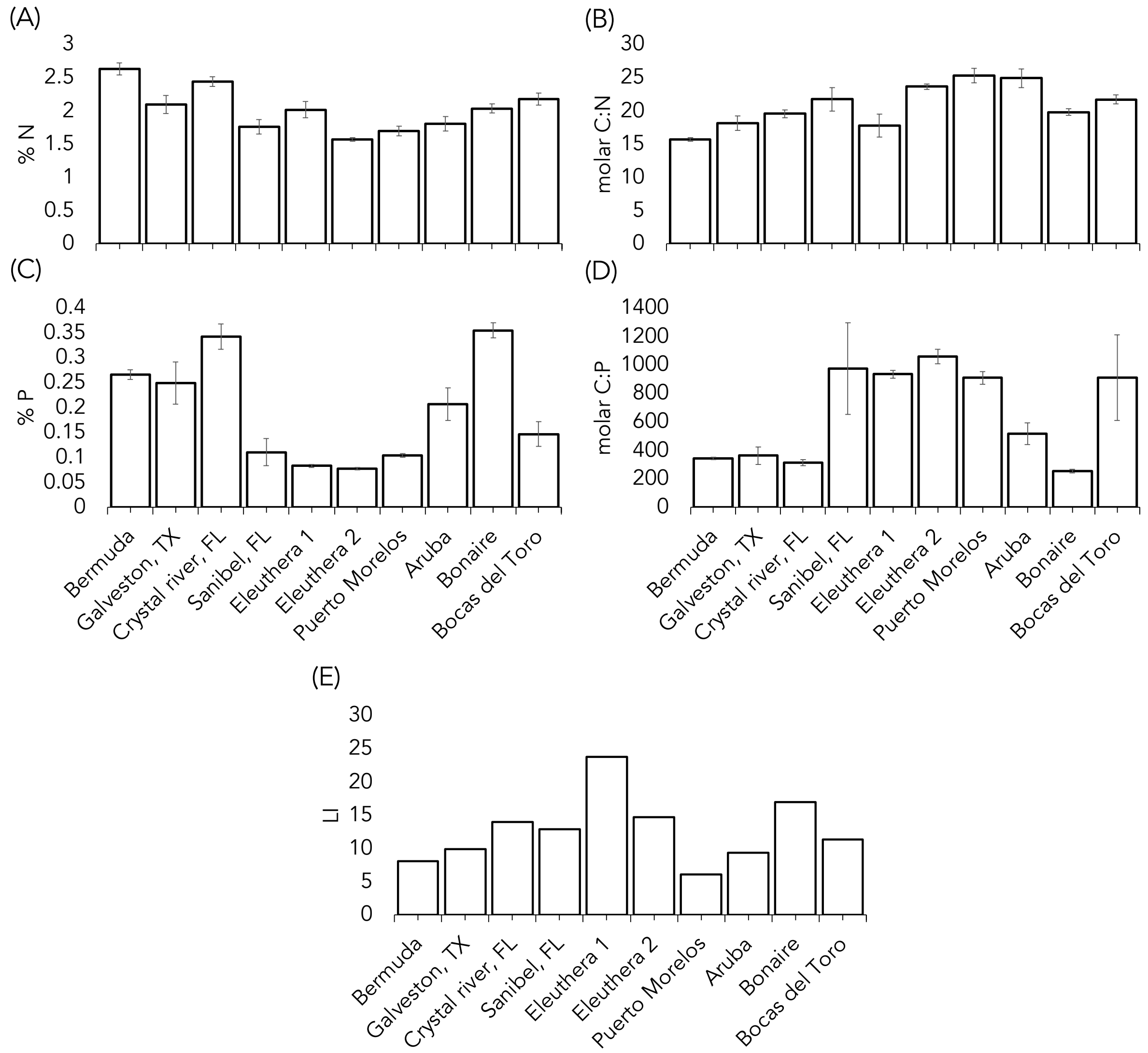


**Supplementary Figure S2**. Trends in leaf (A) %N, (B) molar C:N, (C) %P, (D) molar C:P and (E) Nutrient limitation index across sites in unfertilized plots ± SE.

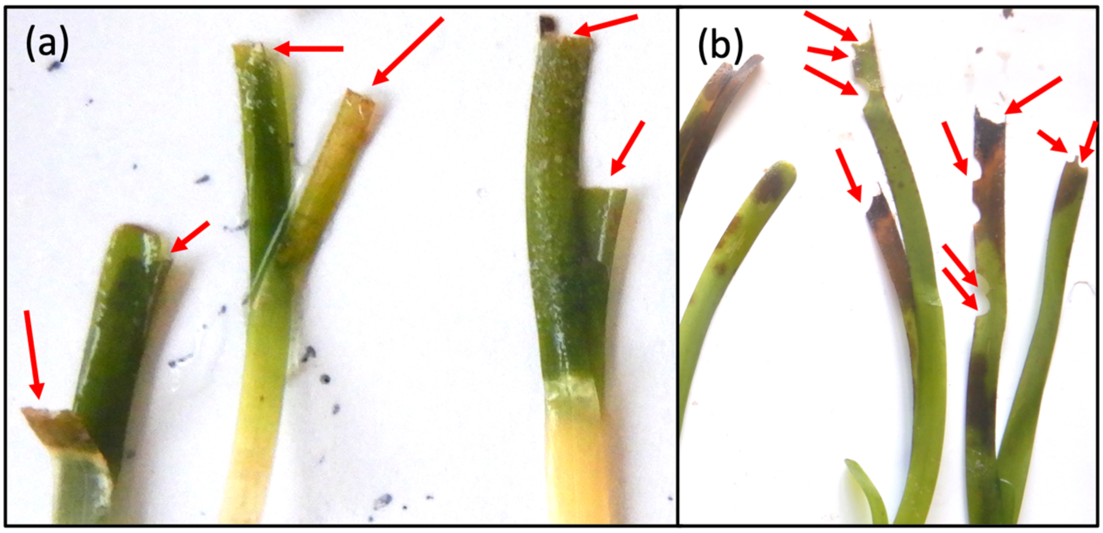


**Supplementary Figure S3**. Differences between green sea turtle grazing and fish grazing (a) *Thalassia testudinum* seagrass shoots that have been grazed by turtles resulting in straight cuts. (b) *T. testudinum* shoots that have been grazed by fish, resulting in crescent shaped bite marks.

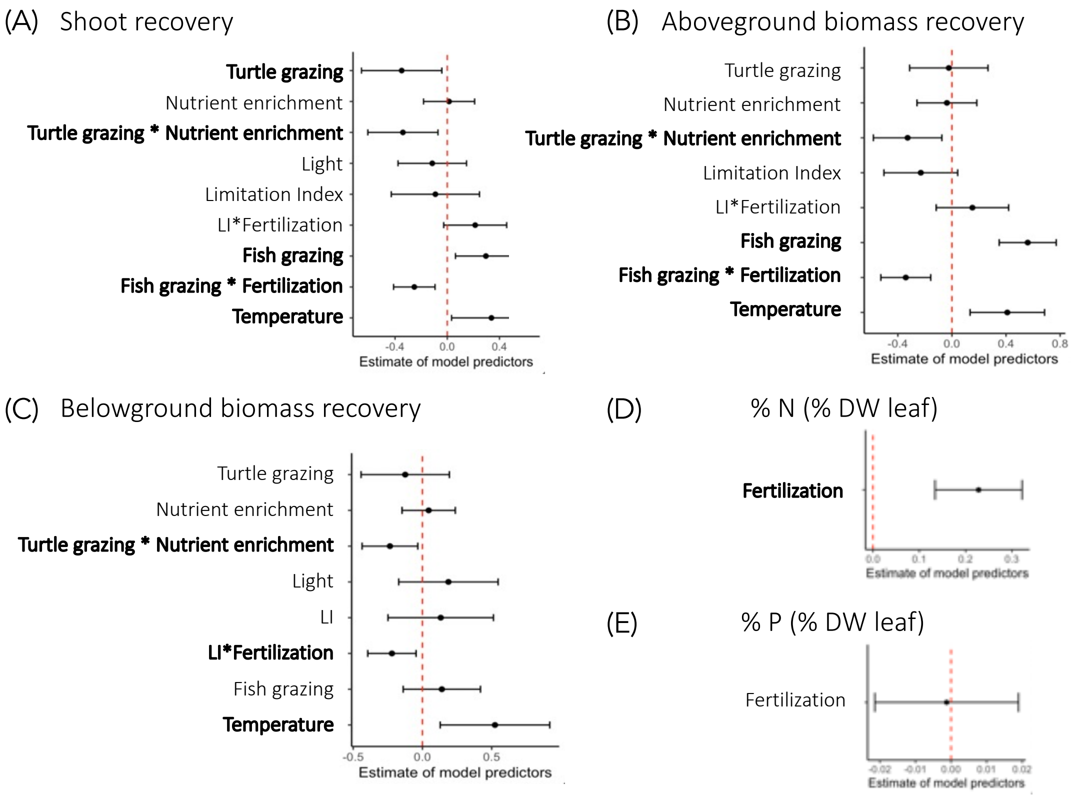


**Supplementary Figure S4.** Standardized coefficient plots displaying model estimates and lines present 95% confidence intervals of the averaged models of (A) shoot recovery, (B) aboveground biomass recovery, (C) belowground biomass recovery, (D) Nitrogen and (E) phosphorus. Significant coefficients are displayed in bold.

**Supplementary Table S1.** Statistical results for averaged generalized linear mixed models testing the impact of fertilization treatments and environmental drivers on seagrass recovery and nutrient content. In this model the effect of seasonality was included instead of the effect of average annual temperature that is presented in the main text. The number of top models (≤ Δ2 AICc) is reported, along with the coefficient estimates and standard errors of the standardized regressors. Seasonality is the SD of temperature among months. Turtle and fish grazing is a grazing index assessed from the leaves. LI is the nutrient limitation index. Light is the yearly average input of light in the system. Nutrient fertilization was simulated by adding both N and P to the water column. Significance codes: ***p<0.0001, **p<0.01, *p<0.05.

| Response | Factor | Estimate | SE | P-value |
|---|---|---|---|---|
| *(A) Shoot recovery* | **Seasonality** | **-0.371** | **0.140** | **0.010*** |
| *(% Shoots* | **Fish grazing** | **0.294** | **0.114** | **0.011*** |
| *compared to* | **Turtle grazing** | **-0.381** | **0.147** | **0.026*** |
| *pre-disturbance)* | **Turtle grazing * Fertilization** | **-0.340** | **0.137** | **0.010**** |
| *(3 top models)* | **Fish grazing * Fertilization** | **-0.257** | **0.081** | **0.002**** |
| | LI * Fertilization | 0.215 | 0.122 | 0.078 |
| | Light | -0.149 | 0.121 | 0.219 |
| | LI | -0.064 | 0.162 | 0.695 |
| | Fertilization | 0.013 | 0.010 | 0.897 |
| *(B) Aboveground* | **Seasonality** | **-0.523** | **0.139** | **0.0002***** |
| *biomass recovery* | **Fish grazing** | **0.398** | **0.119** | **0.0002***** |
| *(% g DW compared* | **Turtle grazing * Fertilization** | **-0.348** | **0.125** | **0.005**** |
| *to pre-disturbance)* | | | | |
| *(3 top models)* | **Fish grazing * Fertilization** | **-0.233** | **0.107** | **0.030*** |
| | LI | -0.181 | 0.135 | 0.178 |
| | LI * Fertilization | 0.149 | 0.133 | 0.263 |
| | Fertilization | -0.023 | 0.110 | 0.834 |
| | Turtle grazing | -0.075 | 0.134 | 0.576 |
| *(C) Belowground* | **Seasonality** | **-0.605** | **0.170** | **0.0009***** |
| *biomass recovery* | **Turtle grazing * Fertilization** | **-0.239** | **0.102** | **0.020*** |
| *(% g DW compared* | | | | |
| *to pre-disturbance)* | **LI * Fertilization** | **-0.219** | **0.089** | **0.014*** |
| *(11 top models)* | Light | 0.135 | 0.168 | 0.421 |
| | Fish grazing | 0.108 | 0.140 | 0.439 |
| | LI | 0.158 | 0.177 | 0.371 |
| | Turtle grazing | -0.179 | 0.158 | 0.257 |
| | Fertilization | 0.037 | 0.010 | 0.708 |

**Supplementary Table S2.** Statistical results for averaged generalized linear mixed models testing the impact of fertilization treatments and environmental drivers on seagrass recovery and nutrient content. In this model the effect of latitude was included instead of the effect of average annual temperature that is presented in the main text. The number of top models (≤ Δ2 AICc) is reported, along with the coefficient estimates and standard errors of the standardized regressors. Seasonality is the SD of temperature among months. Turtle and fish grazing is a grazing index assessed from the leaves. LI is the nutrient limitation index. Light is the yearly average input of light in the system. Nutrient fertilization was simulated by adding both N and P to the water column. Significance codes: ***p<0.0001, **p<0.01, *p<0.05.

| Response | Factor | Estimate | SE | P-value |
|---|---|---|---|---|
| *(A) Shoot recovery (% Shoots compared to pre-disturbance)* | Latitude | -0.249 | 0.188 | 0.185* |
| | **Fish grazing** | **0.282** | **0.126** | **0.027*** |
| | **Turtle grazing** | **-0.344** | **0.170** | **0.043*** |
| | **Turtle grazing * Fertilization** | **-0.311** | **0.128** | **0.015*** |
| *(2 top models)* | **Fish grazing * Fertilization** | **-0.236** | **0.075** | **0.002**** |
| | Fertilization | 0.013 | 0.101 | 0.897 |
| *(B) Aboveground biomass recovery (% g DW compared to pre-disturbance) (10 top models)* | **Latitude** | **-0.489** | **0.182** | **0.007**** |
| | Fish grazing | 0.256 | 0.170 | 0.132 |
| | **Turtle grazing * Fertilization** | **-0.294** | **0.123** | **0.017*** |
| | Fish grazing * Fertilization | -0.193 | 0.108 | 0.074 |
| | Fertilization | -0.021 | 0.113 | 0.856 |
| | Turtle grazing | 0.087 | 0.167 | 0.604 |
| | Light | 0.183 | 0.142 | 0.198 |
| *(C) Belowground biomass recovery (% g DW compared to pre-disturbance) (11 top models)* | Latitude | -0.231 | 0.249 | 0.353 |
| | **Turtle grazing * Fertilization** | **-0.232** | **0.102** | **0.024*** |
| | **LI * Fertilization** | **-0.216** | **0.090** | **0.017*** |
| | Light | 0.277 | 0.241 | 0.250 |
| | Fish grazing | 0.253 | 0.167 | 0.129 |
| | LI | 0.260 | 0.252 | 0.302 |
| | Turtle grazing | -0.236 | 0.180 | 0.190 |
| | Fertilization | 0.029 | 0.099 | 0.768 |

**Supplementary Table S3.** The response of static indicators (measured at one point in time) versus dynamic indicators (% recovery over time) of both shoot density and aboveground biomass to environmental drivers. To analyze this, we compared the unfertilized plots of the recovery experiment to the aboveground biomass and shoot density values as measured at the start of the experiment (N = 5, total of 50 plots) using the same generalized mixed model approach as described in our method section.

| | Response measured | Significant variable | Estimate ± SE | p-value |
|---|---|---|---|---|
| *Static indicators* | | | | |
| | Shoot density (shoots $m^{-2}$) | -- | -- | -- |
| | Aboveground biomass (g DW $m^{-2}$) | | | |
| | | Turtle grazing | **-0.624 ± 0.144** | **0.0003*** ** |
| | | LI | **0.302 ± 0.232** | **0.03*** |
| *Dynamic indicators* | | | | |
| | Shoot recovery (%) | | | |
| | | Temperature | **0.361 ± 0.169** | **0.03** |
| | | Fish grazing | **0.660 ± 0.161** | **0.00004*** ** |
| | Aboveground biomass recovery (%) | | | |
| | | Temperature | **0.612 ± 0.215** | **0.004 ** ** |
| | | Fish grazing | **0.944 ± 0.148** | **<0.001*** ** |
| | | Turtle grazing | 0.442 ± 0.175 | **0.01 *** |
| | | LI | -0.418 ± 0.18 | **0.02 *** |

**Supplementary Table S4.** Table of the experimental sites with the location, start date, end date and duration of the experiment.

| Site | Country | Latitude | Longitude | Start date | End date | Duration (days) |
|---|---|---|---|---|---|---|
| Riddell's Bay | Bermuda | 32°15'49.9"N | 64°49'50.5"W | 14 Sept '18 | 7 Aug '19 | 327 |
| Galveston | Texas, USA | 29°02'41.8"N | 95°10'15.7"W | 2 Oct '18 | 12 Sept '19 | 433 |
| Crystal river | Florida, USA | 28°42'50.4"N | 82°49'08.4"W | 11 Sept '18 | 23 July '19 | 315 |
| Sanibel | Florida, USA | 26°29'48.6"N | 82°09'40.0"W | 15 Sept '18 | 6 Aug '19 | 325 |
| Eleuthera 1 | The Bahamas | 25°27'53.5"N | 76°37'35.8"W | 24 Nov '18 | 10 Nov '19 | 351 |
| Eleuthera 2 | The Bahamas | 25°27'53.7"N | 76°37'35.3"W | 28 Nov '18 | 6 Nov '19 | 343 |
| Puerto Morelos | Mexico | 20°52'04.5"N | 86°51'35.4"W | 19 Sept '18 | 1 Aug '19 | 316 |
| Barcadera Bay | Aruba | 12°28'33.2"N | 69°59'24.0"W | 16 July '18 | 17 Apr '19 | 305 |
| Lac Bay | Bonaire, NL | 12°06'44.3"N | 68°13'42.0"W | 12 Sept '18 | 16 Oct '19 | 399 |
| Bocas del Toro | Panama | 9°21'05.8"N | 82°15'27.8"W | 26 Sept '18 | 16 July '19 | 293 |

**Supplementary Table S5.** Results of the regression analysis testing the linear average shoot recovery in unfertilized plots over time

| Location | Estimate | SE | t value | P value | $R^2$ |
|---|---|---|---|---|---|
| Riddell's bay, Bermuda | 0.002 | 0.0008 | 2.587 | 0.036* | 0.49 |
| Galveston, TX, USA | 0.003 | 0.001 | 2.561 | 0.043* | 0.52 |
| Crystal river, FL, USA | 0.014 | 0.006 | 2.222 | 0.048* | 0.68 |
| Sanibel, FL, USA | 0.006 | 0.003 | 2.323 | 0.033* | 0.29 |
| Eleuthera 1, Bahamas | 0.116 | 0.010 | 11.30 | 0.011* | 0.99 |
| Eleuthera 2, Bahamas | 0.009 | 0.004 | 2.359 | 0.078 | 0.58 |
| Puerto Morelos, Mexico | 0.014 | 0.001 | 9.541 | 0.0000006*** | 0.88 |
| Barcadera bay, Aruba | 0.005 | 0.0007 | 7.029 | 0.00001*** | 0.80 |
| Lac bay, Bonaire | 0.026 | 0.003 | 8.442 | 0.00006*** | 0.91 |
| Bocas del Toro, Panama | 0.010 | 0.002 | 5.990 | 0.0002*** | 0.80 |

**Supplementary Table S6.** Pearson correlation matrix showing correlation coefficients between candidate environmental drivers of recovery rates. Latitude and seasonality were excluded from the multi-model inference because r > 0.50 with temperature.

| | Fish grazing | Avg temp | Latitude | LI | Turtle grazing | Seasonality |
|---|---|---|---|---|---|---|
| Turtle grazing | | | | | | 0 |
| LI | | | | | 0.4 | -0.2 |
| Latitude | | | | 0 | 0.4 | 0.7 |
| Avg temp | | | -0.7 | 0.3 | -0.2 | -0.9 |
| Fish grazing | | 0.3 | -0.4 | 0.3 | -0.1 | -0.4 |
| Light | 0.3 | 0.2 | -0.1 | 0.4 | 0 | -0.3 |

**Supplementary Table S7.** Model selection table of the top models explaining shoot recovery. Presented are AICc values with the delta deviation from the top model, and the $R^2$ variation that is explained by the models both as conditional and marginal $R^2$

| Response | Model parameters | AICc | delta | conditional $R^2$ | marginal $R^2$ |
|---|---|---|---|---|---|
| Shoot recovery | 1: fertilization + temperature + fish grazing + turtle grazing, + fertilization* turtle grazing + fertilization *fish grazing | 779.3 | 0.00 | 0.501 | 0.408 |
| | 2: fertilization + temperature + fish grazing + turtle grazing + LI + fertilization * turtle grazing + fertilization * fish grazing + fertilization * LI | 780.8 | 1.51 | 0.527 | 0.438 |
| | 3: fertilization + fish grazing + turtle grazing + fertilization * turtle grazing + fertilization * fish grazing | 781.0 | 1.70 | 0.461 | 0.290 |
| | 4. Fertilization + temperature + fish grazing + turtle grazing, + fertilization* turtle grazing + fertilization *fish grazing + Light | 781.2 | 1.82 | 0.495 | 0.419 |
| Aboveground biomass recovery | 1: fertilization + temperature + fish grazing + turtle grazing + LI + fertilization * turtle grazing + fertilization * fish grazing | 825.2 | 0.00 | | 0.423 |
| | 2: fertilization + temperature + fish grazing + turtle grazing + fertilization * turtle grazing + fertilization * fish grazing | 825.5 | 0.30 | | 0.396 |
| | 3: fertilization + temperature + fish grazing + turtle grazing + LI + fertilization * LI | 826.7 | 1.49 | | 0.431 |
| Belowground biomass recovery | 1: temperature | 888.8 | 0.00 | 0.460 | 0.251 |
| | 2: fertilization + temperature + LI + fertilization * turtle grazing + fertilization * fish grazing + fertilization * LI | 888.9 | 0.17 | 0.534 | 0.340 |
| | 3: temperature + Light | 889.9 | 1.14 | 0.454 | 0.281 |
| | 4: temperature + fish grazing | 890.0 | 1.27 | 0.470 | 0.260 |
| | 5: fertilization + temperature + turtle grazing +fertilization * turtle grazing | 890.1 | 1.37 | 0.513 | 0.293 |
| | 6: fertilization + temperature + fish grazing + LI + fertilization * LI | 890.4 | 1.59 | 0.505 | 0.311 |
| | 7: temperature + LI | 890.4 | 2.63 | 0.457 | 0.267 |
| | 8: fertilization + temperature + LI + fertilization * LI + Light | 890.5 | 1.72 | 0.489 | 0.328 |
| | 9: fertilization + temperature + turtle grazing + LI + + fertilization * LI + | 890.6 | 1.84 | 0.493 | 0.321 |
| | 10: temperature + turtle grazing | 890.9 | 1.99 | 0.467 | 0.255 |

**Supplementary Equation S1.** Model structure

*y* ~ fertilization + turtle grazing + turtle grazing x fertilization + fish grazing + fish grazing x fertilization + temperature + limitation_index + limitation_index x fertilization + light + (1|site)
